# Inferring propensity amongst lung and breast carcinomas via overlapped gene expression profiles

**DOI:** 10.1101/178558

**Authors:** Rajni Jaiswal, Sabin Dhakal, Shaurya Jauhari

## Abstract

Reconstruction of biological networks for topological analyses helps in correlation identification between various types of biomarkers. These networks have been vital components of System Biology in present era. Genes are the basic physical and structural unit of heredity. Genes act as instructions to make molecules called proteins. Alterations in the normal sequence of these genes are the root cause of various diseases and cancer is the prominent example disease caused by gene alteration or mutation. These slight alterations can be detected by microarray analysis. The high throughput data obtained by microarray experiments aid scientists in reconstructing cancer specific gene regulatory networks. The purpose of experiment performed is to find out the overlapping of the gene expression profiles of breast and lung cancer data, so that the common hub genes can be sifted and utilized as drug targets which could be used for the treatment of diseased conditions. In this study, first the differentially expressed genes have been identified (lung cancer and breast cancer), followed by a filtration approach and most significant genes are chosen using paired t-test and gene regulatory network construction. The obtained result has been checked and validated with the available databases and literature.

## 1. INTRODUCTION

Gene Regulatory Network (GRN) is a systemic biological network that helps to understand the interaction between two genes in the form of a pictorial representation. In the diagram, the nodes represent the genes studied whereas the edges represent their regulatory interactions. GRN helps to understand the activation and inhibition of specific genes by their counterparts present in the same cell [2, 6].

The disease in question is Cancer, which is also known as a 'malignant neoplasm'. It entails uncontrolled and unregulated cell growth. The cancerous cells (oncogenes) have a tendency to divide and multiply uncontrollably, leading to the formation of malignant tumors and infesting the nearby regions of the body through lymphatic system. This swarm may lead to the formation of a new (secondary) tumor, distant to the original (primary) tumor. Chemotherapy, radiotherapy and surgery could be the possible means for the diagnosis and treatment of cancer. However, these methods of treatment often have deleterious effect in healthy cells and tissues. Therefore, identification of molecular markers of cancers could be an alternative approach to diagnose the human cancer and might be useful for development of novel therapies [1]. Although, various significant genes responsible for the genesis of different tumors have been unraveled but fundamental molecular interactions are still unclear and remains a challenge for the researchers [1]. GRNs have proved to be a very useful tool to explain complex dependencies between the key developmental transcription factors, their target genes and regulators. In this work, information-theoretic approach called *mutual information* of diseased and normal condition has been used to compute regulatory relationships between gene-pairs. We applied this approach to reconstruct GRN of lung and breast cancer which are two leading cause of cancer mortalities world-wide.

Among U.S. women, breast cancer is commonly prevalent and is revered second leading cause of death after lung cancer. In 2014, an estimated 232,670 new cases of invasive breast cancer were expected to be diagnosed in women in the U.S., along with 62,570 new cases of non-invasive (in situ) breast cancer [13]. Also metastatic breast cancer, which is spreadable to different tissues including lung, leads to many cases of lung cancer. It has been found that genes responsible for breast cancer metastasis to lungs and are clinically correlated in development of lung metastasis when expressed in primary breast cancer [10]. Radio therapy is a conventional treatment process for the regionally advanced breast cancer; however, it was found that breast cancer radiation therapy increased the risk of lung cancer especially in cigarette smokers [11]. These findings signify strong correlation between breast and lung cancer, and intuitively we can deduce that there might be mutual genes which are responsible for both disease states. Some genes could be non-functional in their own expression levels however could have some expression in interaction with another gene or could lead to the expression of another gene as a part of gene network. Identification of such network could be vital for the targeted therapy of the cancer condition if we are able to find out some common gene in both the cancer conditions.

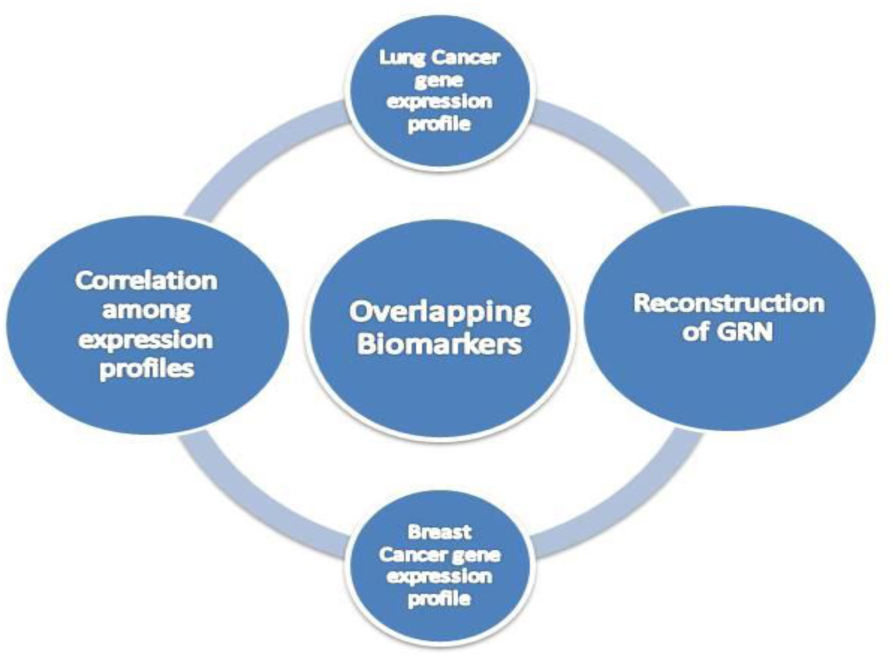

## 2. METHODS

i. Dataset pre-processing.
ii. Analysis of most significant genes.
iii. Correlation identification between gene pairs.
iv. Reconstruction of Gene Regulatory Network (GRN).
v. Identification of Hub genes and common genes.
vi. Biological validation.

### 2.1. Dataset pre-processing

The dataset preprocessing includes normalization, removal of noisy data and duplicates from dataset. Normalization is a process in which data attributes within a data model are organized to reduce or even eliminate the redundancy. As each of the GDS file is already internally normalized as part of the uploading requirement to GEO, intra-dataset normalization was not necessary. We did data pre-processing to handle missing gene names, noisy data and the duplicates were removed by using function:

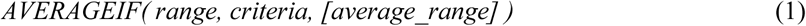

### 2.2. Analysis of most significant genes

In this step, we tried to analyse the genes that show high expression in diseased condition. T-test is used to determine if two sets of data are significantly different from one another. The t-test can be computed as follows:

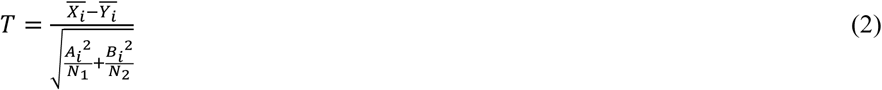

where X_i_ and Y_i_ are means of the test and control samples, 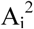 and 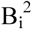 are variances of test and control and N1 and N2 are sizes of test and control, respectively, of i which is gene expression profile. During the test we considered two sets of the data as unequal variance, *p-value* which are <=0.001 only considered to find out the most significant genes.

### 2.3. Correlation identification between gene pairs

Finding of co-relationship between the gene pairs, we applied a statistical method i.e Pearson Correlation Coefficient in this study. The linear associations vary between ± 1, where r = 1 means a perfect positive correlation and the value r = -1 mean perfect negative correlation. For r = 0, variables are independent.

The formula for Pearson Correlation Coefficient (R) is:

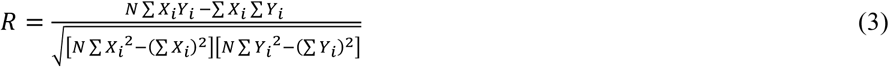

After computing pair-wise correlation coefficient, we selected those coefficients whose absolute value is above a threshold (0.75 in case of breast cancer and 0.94 in case of lung cancer). The strategy for this paper is to focus on highly connected genes. We computed the pair-wise correlation coefficient among genes in both Breast Cancer and Lung Cancer and observed mostly positive and few negative in breast cancer where in lung cancer, all connections are positive.

### 2.4. Reconstruction of Gene Regulatory Network (GRN)

In this step, biological networks for both diseases were constructed using Cytoscape tool. Cytoscape is an open source software platform for visualizing complex networks and integrating these with any type of attribute data.

## 3. RESULTS AND DISCUSSIONS

In the present manuscript, microarray data set of breast cancer and lung cancer has been taken for gene regulatory network construction, topological analysis and deciphering common interaction between the two types of cancer. The full dataset was downloaded from NCBI (National Centre for Biotechnology Information) within the sub search level of GEO Datasets (<u>http://www.ncbi.nlm.nih.gov/sites/GDSbrowser/</u>)[30] which consists of following microarray data:

Breast cancer: 21766 genes, 42 samples, dataset _ reference _ series = GSE20437

Lung cancer: 22215 genes, 192 samples, dataset _ reference _ series = GSE4115

A series of steps were performed in order to form the gene regulatory network. First stage was pre-filtering of data to segregate out noisy data such as genes with unknown names or the ones which do not exist in the NCBI gene library and also duplicates were removed by taking average of the repeated genes. The number of genes that remained

Breast cancer: 13652 (62.7% genes were remaining.)

Lung cancer: 13056 (58.8% genes were remaining.)

To find out the most significant genes, a single step filtration with t-test statistics was performed and those genes having p value <=0.001 were the only ones considered. The number of genes remaining after filtering out the non-significant genes by t-test is as follows:

Breast cancer: 50 (0.22% genes were remaining.) Lung cancer: 639 (2.9% genes were remaining.)

Further, Pearson correlation was applied in order to measure the strength of pair-wise correlation between the extracted genes of each type of cancer. From the resultant genes with weak Pearson correlation coefficient, the respective gene pairs have been omitted. The absolute value for correlation which has been deemed significant is 0.75 and 0.94 for breast cancer and lung cancer, correspondingly. The strong correlation involved 45 relations in Breast cancer and 65 in Lung cancer. Finally, extracted genes were validated with available biological databases and literature. Table 1 below shows the interaction of gene pairs marking the relation as activating (+) or repressing (-). The positive (+) correlation shows activation and negative value (-) shows repression (inhibiting).

Breast cancer: Out of 45 regulatory relations, 37 are activators. 8 are repressors.

**Table 1:** List of genes found to be involved in Breast cancer.

| Source | Target | activator (+),<br>repressor (-) |
| --- | --- | --- |
| *BTG2* | *ATF3* | + |
| *DDX39A* | *CAPZB* | + |
| *FOS* | *DUSP1* | + |
| *FOSB* | *DUSP1* | + |
|  | *FOS* | + |
| *H3F3B* | *DUSP1* | + |
| *IER2* | *ATF3* | + |
|  | *BTG2* | + |
|  | *DUSP1* | + |
| *IFRD2* | *C19orf10* | + |
|  | *DDX39A* | + |
| *ILF3* | *ACTR10* | - |
|  | *DDX39A* | + |
|  | *ENDOD1* | - |
|  | *IFRD2* | + |
| *JUN* | *ATF3* | + |
|  | *BTG2* | + |
|  | *DUSP1* | + |
|  | *IER2* | + |
| *KLF11* | *CAPZB* | - |
|  | *ENDOD1* | + |
| *PCYT2* | *DDX39A* | + |
|  | *IFRD2* | + |
|  | *ILF3* | + |
| *PDIA4* | *C19orf10* | + |
| *RNF139* | BTG2 | + |
|  | DDX39A | - |
|  | ILF3 | - |
| *RORA* | ENDOD1 | + |
|  | KLF11 | + |
| *SNRK* | BTG2 | + |
|  | ENDOD1 | + |
|  | ILF3 | - |
|  | PCYT2 | - |
|  | RNF139 | + |
| *WIPF2* | ENDOD1 | + |
| *YPEL5* | ATF3 | + |
|  | BTG2 | + |
|  | ILF3 | - |
|  | RNF139 | + |
|  | SNRK | + |
| *YTHDC1* | RARS | + |
|  | RNF139 | + |
|  | SNRK | + |
| *ZNF318* | RNF139 | + |

Lung Cancer: Out of 65 regulations R >=0.94 all are activators.

**Table 2.**
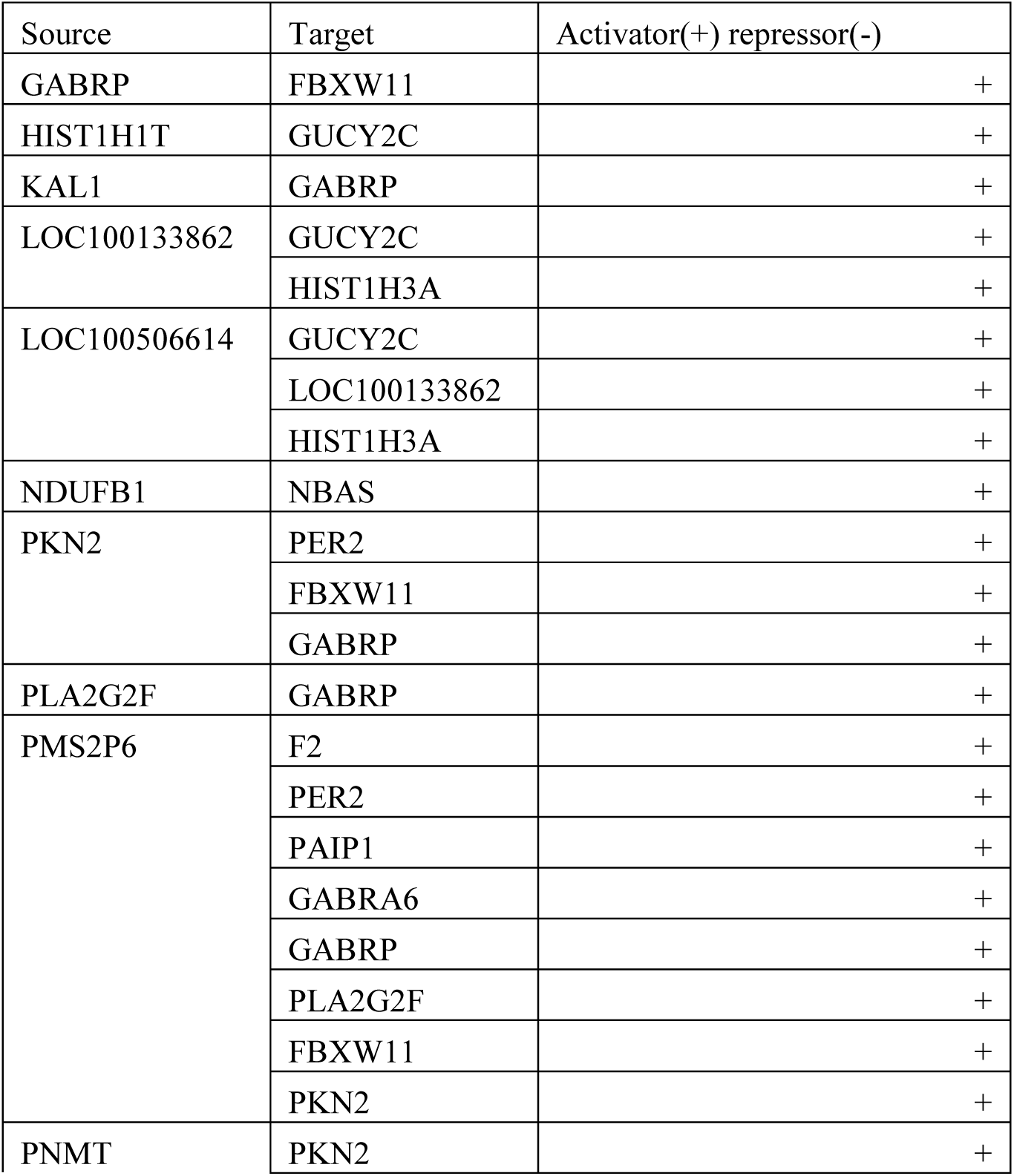
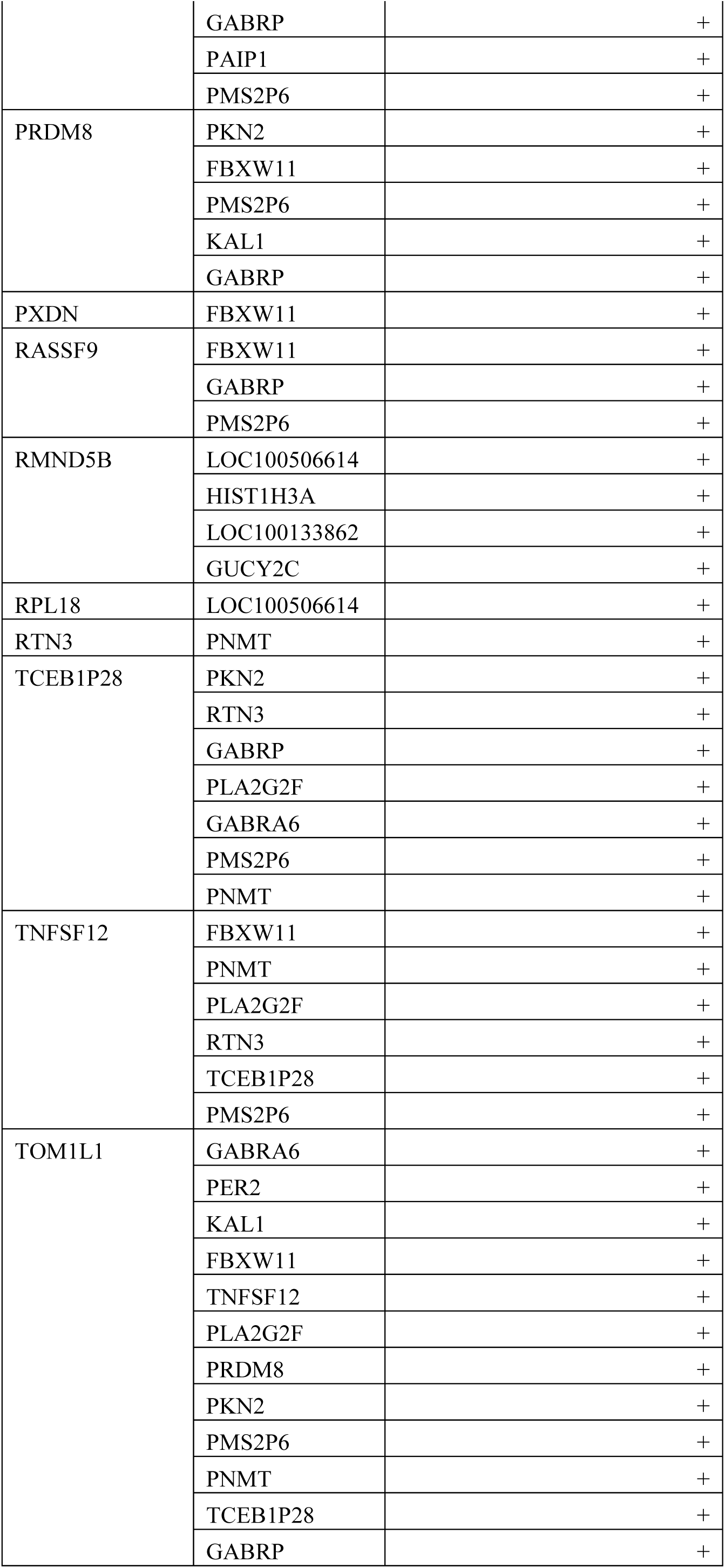
List of genes found to be involved in Lung cancer

**Figure 1:**
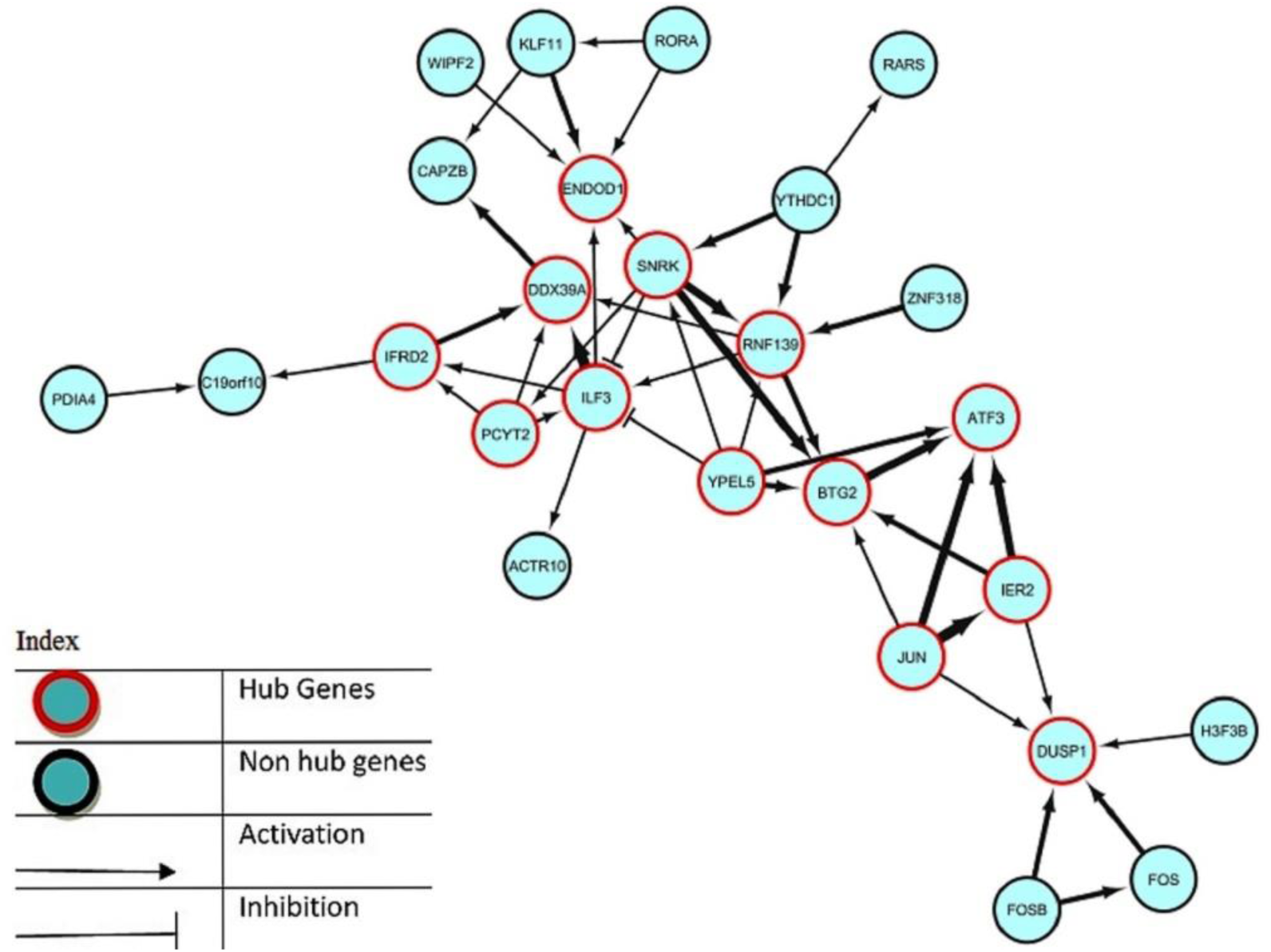
Inferred gene regulatory network of 26 genes and 45 regulatory relations of Breast Cancer using proposed methodology. The finding shows that the genes in red outline circles are involved as hub genes.

**Table 3:**
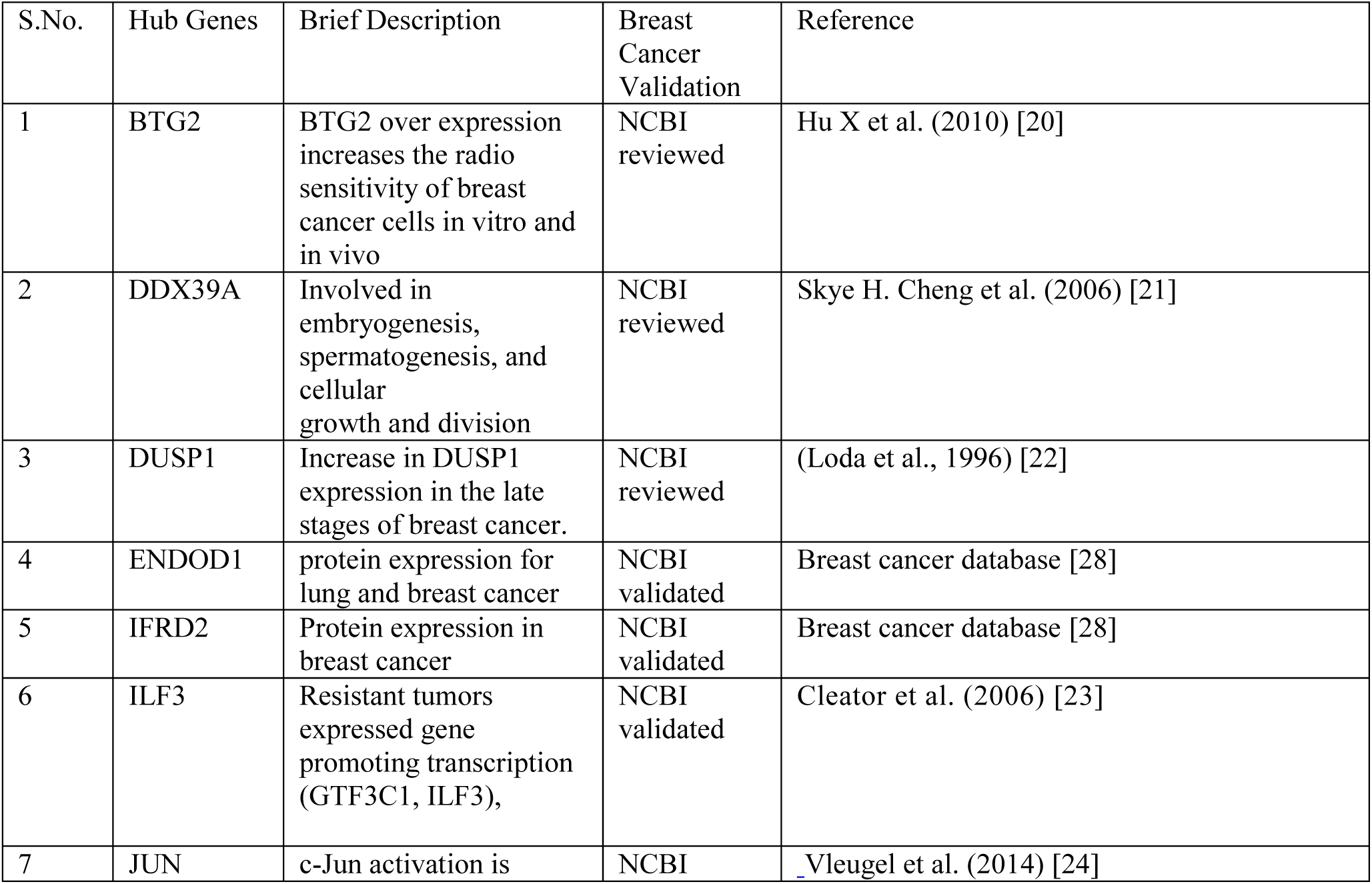
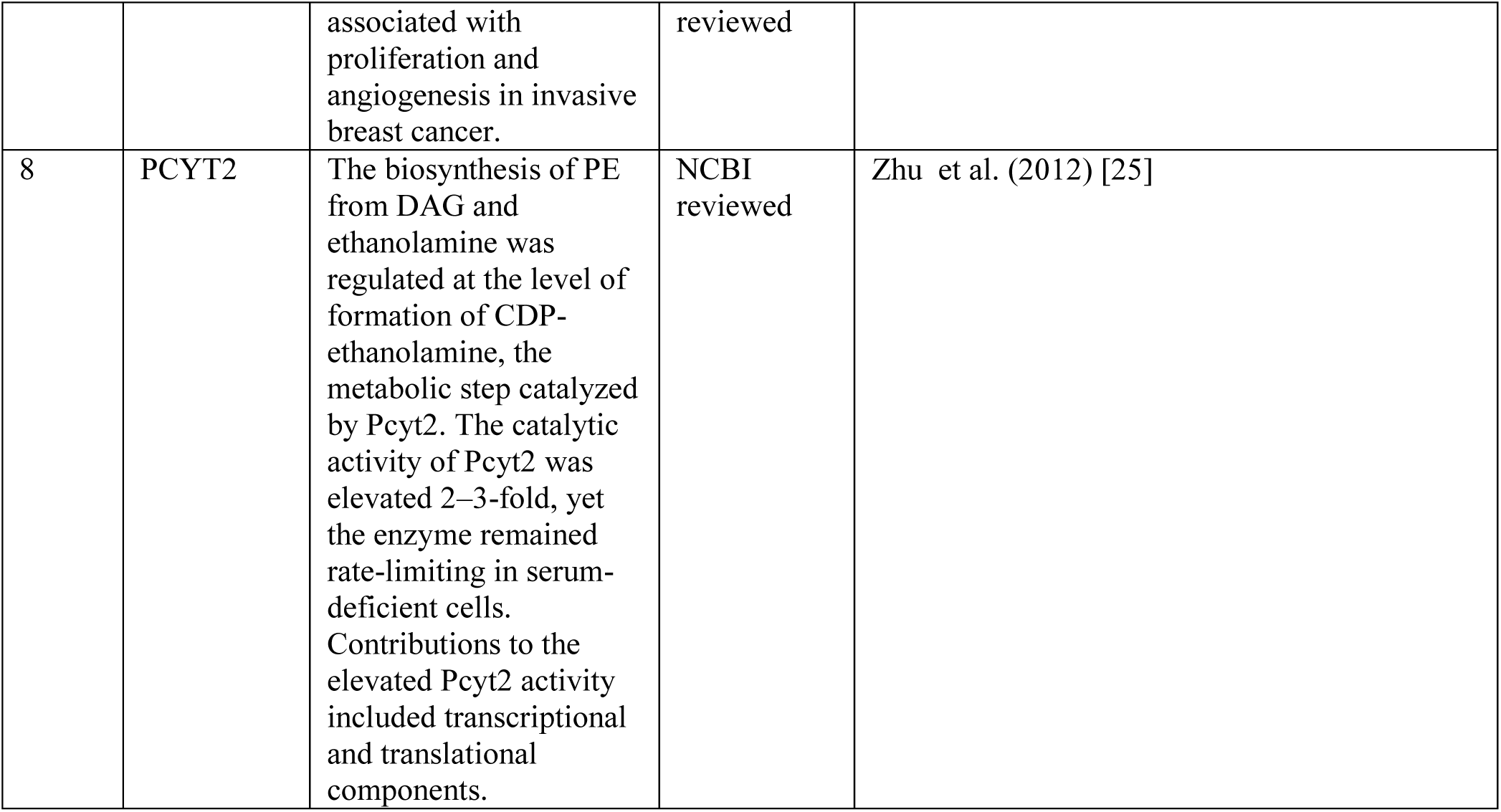
*Validated list of genes involved in Breast Cancer*

**Figure 2:**
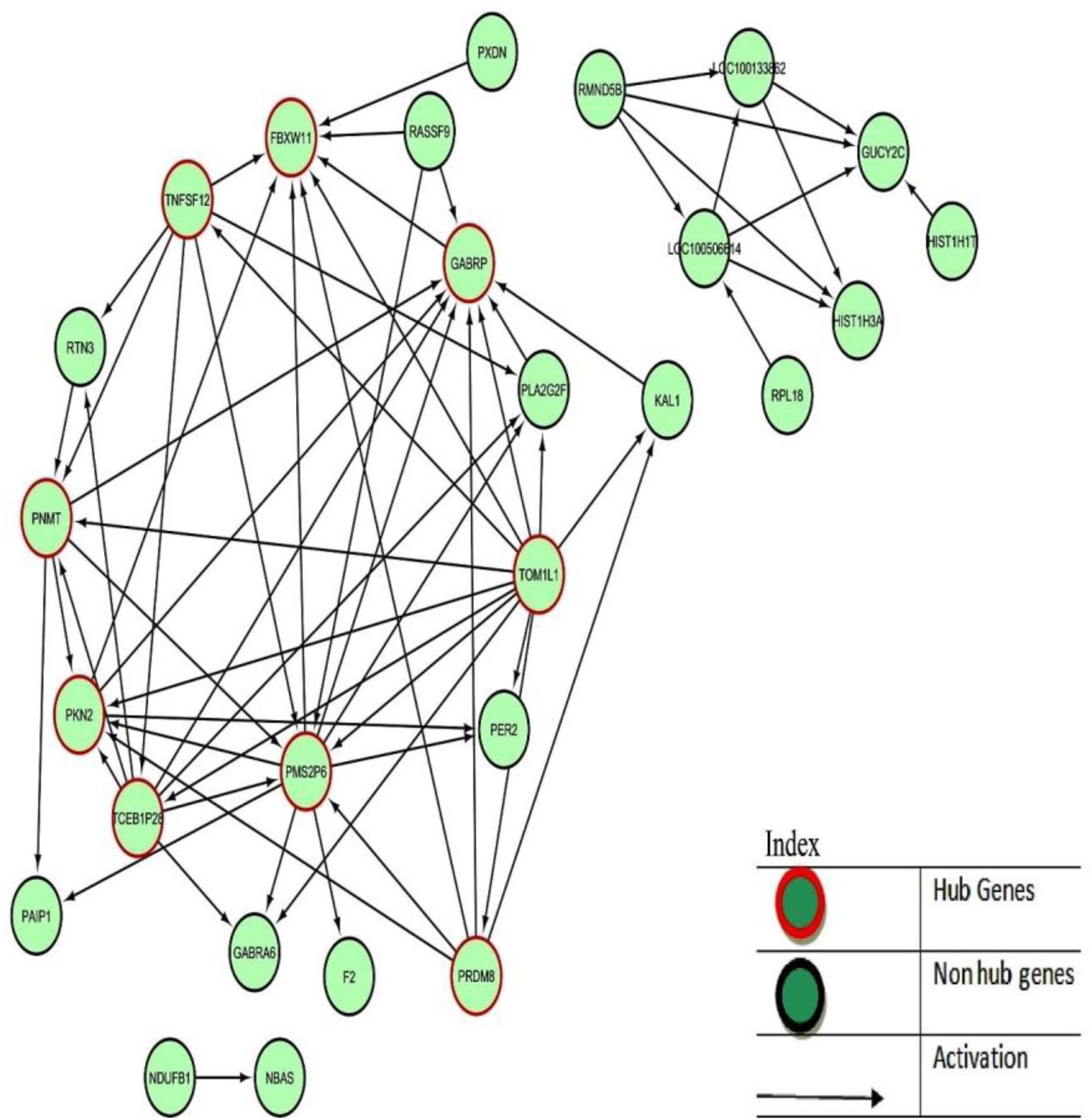
Inferred gene regulatory Network of 27 genes and 65 regulatory relations in Lung Cancer using proposed methodology. Genes in red outline circles are participating as hub Genes.

**Table 4.** Validated List of genes involved in Lung Cancer.

| S.No | Gene Name | Brief Description | References |
| --- | --- | --- | --- |
| 1. | GABRP | Over-expressed in various tumors, including Lung cancer | Human Protein atlas experimental results. [32] |
| 2. | FBXW11 | FBXW11/BTRC control over RAPGEF2 protein levels was observed in H2228 lung adenocarcinoma cells. | Tai Young Kim et al. 2015 [26] |
| 3. | TNFSF12 | Weak inducer of apoptosis in some cell types. Mediates NF-kappa-B activation. Promotes angiogenesis and the proliferation of endothelial cells. | NCBI [30] |
| 4. | TOM1L1 | Probable adapter protein involved in signaling pathways. Interacts with the SH2 and SH3 domains of various signaling proteins when it is phosphorylated. May promote FYN activation, possibly by disrupting intramolecular SH3-dependent interactions | Gene Mania Database [33] |
| 5. | PRDM8 | Highly expressed in various cancer including lung, breast, prostate, liver. | Human protein atlas data. [32] |
| 6. | PKN2 | PKN2 identified in extracellular vesicles derived from the tissue/cell of Lung cancer and many other cancers | NCBI [30] |
| 7. | PMS2P6 | Genetic variation in the DNA repair gene is predictive of outcome in lung cancer. | Athena Matakidou et al. (2007) [27] |

### 3.1 Common expression profiles in Breast Cancer and Lung Cancer

We have identified highly expressed common gene profiles with p value <=0.001 and <=0.01. This strategy helps us to find out the involvement of few genes which show high expression in both cancer types. Study of these genes can further help in understanding of their function and approach to the targeted therapy for the patients suffering from breast cancer and having high risk of getting lung cancer.

The common genes and their corresponding p-value given below:

**Table 5.** Table showing the common highly expressed genes on breast cancer and lung cancer and their corresponding p-value of lung cancer (<=0.001).

| p-value in Lung cancer ($\leq 0.001$) | Common Highly expressed genes | p-value in breast cancer ($\leq 0.001$) |
| --- | --- | --- |
| 0.000198 | ATP6V1B2 | 0.000325 |
| 0.000428 | PCYT2 | 0.000891 |
| 0.000131 | KLF11 | 7.85E-05 |

**Table 6:** Table showing the common genes on breast and lung cancer of p value <=0.01.

| p-value in lung cancer | Common genes | p-value in breast cancer |
| --- | --- | --- |
| 0.000197849 | ATP6V1B2 | 0.000325295 |
| 4.35E-05 | BAZ1B | 0.003824184 |
| 0.000471294 | DECR2 | 0.002539938 |
| 0.000780486 | HDAC3 | 0.001499691 |
| 1.84E-05 | HEXB | 0.004434946 |
| 0.000209637 | IER3 | 0.003053135 |
| 0.000887335 | IRF6 | 0.003370621 |
| 0.000130787 | KLF11 | 7.85E-05 |
| 0.000373077 | LARS2 | 0.004304204 |
| 0.00017249 | LOC100508591 | 0.00839416 |
| 0.000460253 | LRRC31 | 0.005627496 |
| 0.000998246 | LZTS3 | 0.001050066 |
| 0.000870628 | NEK11 | 0.009949024 |
| 1.09E-05 | PAQR6 | 0.004034503 |
| 0.000427581 | PCYT2 | 0.00089129 |
| 0.000318592 | RPS11 | 0.006368626 |
| 2.50E-06 | RTN3 | 0.005591664 |
| 1.77E-08 | SET | 0.007373827 |
| 1.77E-05 | SSFA2 | 0.008010013 |
| 0.000984356 | TNFSF12-TNFSF13 | 0.003453627 |

### 3.2 Biological Validation of common highly expressed genes

**Table 7:**
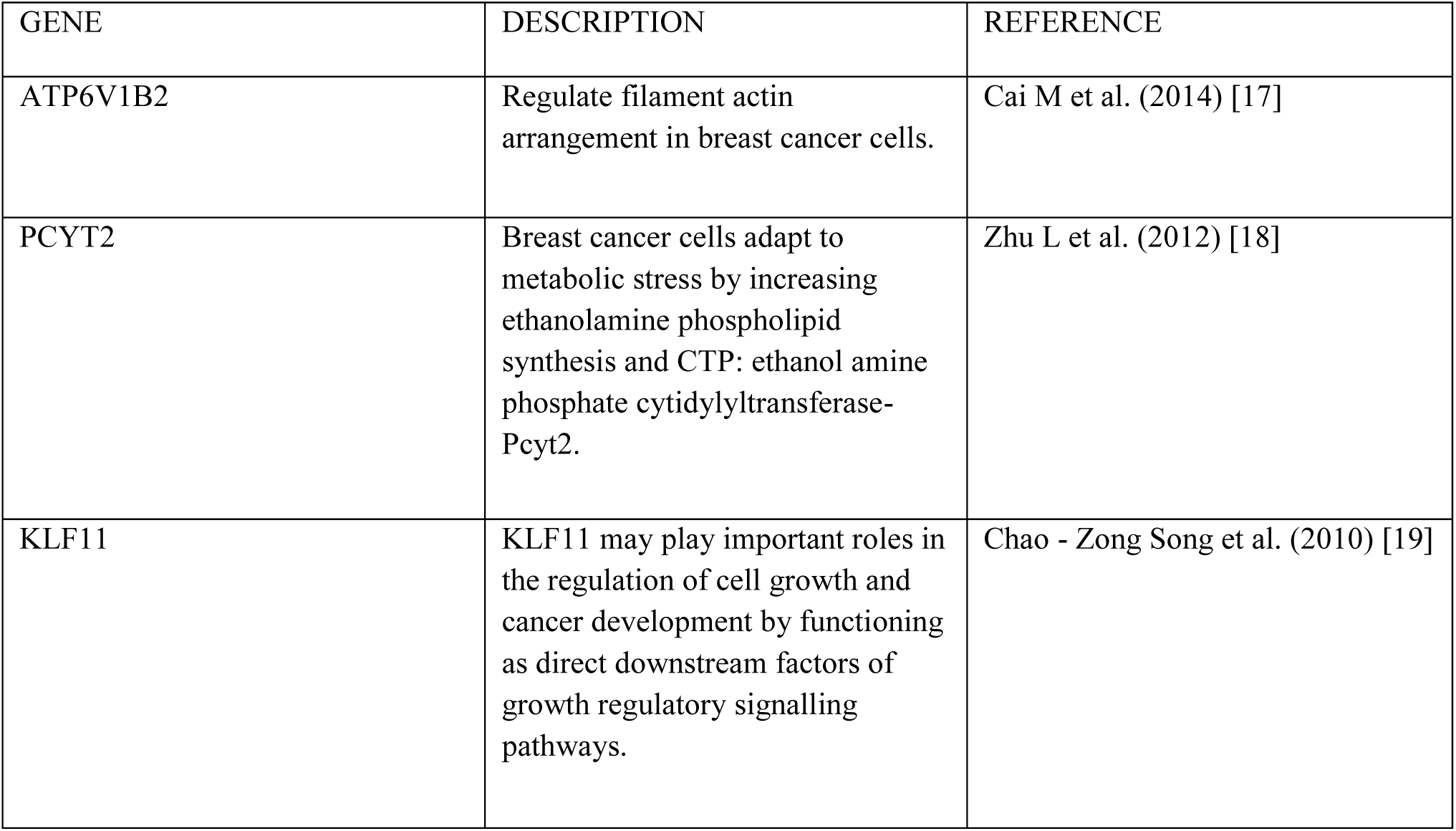
Table showing the validation of gene with various databases

| GENE | DESCRIPTION | REFERENCE |
| --- | --- | --- |
| ATP6V1B2 | Regulate filament actin arrangement in breast cancer cells. | Cai M et al. (2014) [17] |
| PCYT2 | Breast cancer cells adapt to metabolic stress by increasing ethanolamine phospholipid synthesis and CTP: ethanol amine phosphate cytidyltransferase-Pcyt2. | Zhu L et al. (2012) [18] |
| KLF11 | KLF11 may play important roles in the regulation of cell growth and cancer development by functioning as direct downstream factors of growth regulatory signalling pathways. | Chao - Zong Song et al. (2010) [19] |

## 4. CONCLUSION

Cancer specific Gene Regulatory Networks provide key information for identification of cancerous genes and their pathways. A directed regulatory network is capable of revealing interactions among genes more reasonably and also capturing cause-effect relations between gene-pairs. This report is a simple statistical approach to extract differentially expressed genes, finding correlations between gene-pairs for the reconstruction of gene regulatory networks underlying different specific disease conditions like breast cancer and lung cancer that assist the interpretability of the network.

From the analysis of the constructed network, we observed that some genes are working as hub genes including SNRK, ILF3, RNF139, BTG2, and ENDOD1. Among them, BTG2 is highly linked in breast cancer. The further study of these genes in the metastasis level of breast cancer helps in exploration of the association of lung cancer by finding lung cancer tumorigenesis with breast cancer metastasis.

Similarly, GABRP, PNMT, FBXW11, PKN2, PMS2P6, TOM1L1, TCEB1P28, PRDM8, TNFSF12 are hub genes. Among them TOM1L1 and PMS2P6 are found to be highly linked in lung cancer.

The efficiency of the results is explained by the gene validation done thorough various literature and database review.

As per the GRN we did not find any hub genes which are commonly present in both the cancerous condition with significant correlation coefficient. However, there are common genes: ATP6V1B2, PCYT2, KLF11 which are highly expressed in both the cancerous condition.

Similarly, BAZ1B, DECR2, HDAC3, HEXB, IER3, IRF6, LARS2, PAQR6 and other genes listed in Table 4 are found to have significant levels of expression in both the cancer conditions. Our findings help to reveal common molecular interactions in breast and lung cancer studies and provide new insights in targeted cancer diagnostics, prognostics and therapy in the population who are highly susceptible to the breast cancer and have likely chance of developing lung cancer. Moreover, further exploration of the commonly expressed genes helps the medicinal chemistry discipline to develop the common therapeutic approach to control both cancer types which is more challenging to the scientists in present world.

The proposed approach along with the more extensive computational biology can also be used to investigate other cancer specific gene regulatory network like colon cancer, blood cancer, throat cancer and others. As an attempt to scale our study, we aim to construct regulatory networks for other types of cancer from microarray data and try to establish common root to treat various cancer types.

## 5. LIST OF ABBREVIATIONS

GRN: Gene Regulatory Network
GDS: Graphic Data System File
GEO: Gene Expression Omnibus
NCBI: National Centre for Biotechnology Information
GSE: Genomic Spatial Event database
SNRK: Sucrose Nonfermenting (SNF)-Related Kinase
ILF3: Interleukin enhancer-binding factor 3
RNF139: Ring Finger Protein 139
BTG2: B-cell translocation gene
2 ENDOD1: Endonuclease Domain Containing 1
GABRP: Gamma-Aminobutyric Acid Type A Receptor Pi Subunit
PNMT: Phenylethanolamine N-methyltransferase
FBXW11: F-Box And WD Repeat Domain Containing 11
PKN2: Protein Kinase N2
PMS2P6: PMS1 homolog 2, mismatch repair system component pseudogene 6
TOM1L1: Target Of Myb1 Like 1 Membrane Protein
TCEB1P28: Transcription Elongation Factor B Subunit 1 Pseudogene 28
PRDM8: PR/SET Domain 8
TNFSF12: Tumor Necrosis Factor Superfamily Member 12)
ATP6V1B2: ATPase H+ Transporting V1 Subunit B2
PCYT2: Phosphate cytidylyl Transferase 2
KLF11: Kruppel Like Factor 11
BAZ1B: Bromodomain Adjacent To Zinc Finger Domain 1B
DECR2: 2,4-Dienoyl-CoA Reductase 2
HDAC3: Histone Deacetylase 3
HEXB: Hexosaminidase Subunit Beta
IER3: Immediate Early Response 3
IRF6: Interferon Regulatory Factor 6
LARS2: Leucyl-TRNA Synthetase 2
PAQR6: Progestin And AdipoQ Receptor Family Member 6

## 7. ACKNOWLEDGEMENT

The authors would like to thank Mr. Mukul Varsney, Assistant Professor, Department of Computer Science Engineering, Sharda University for his support in the work.

